# PyDESeq2: a python package for bulk RNA-seq differential expression analysis

**DOI:** 10.1101/2022.12.14.520412

**Authors:** Boris Muzellec, Maria Teleńczuk, Vincent Cabeli, Mathieu Andreux

## Abstract

**Summary:** We present PyDESeq2, a python implementation of the DESeq2 workflow for differential expression analysis on bulk RNA-seq data. This implementation achieves better precision, allows speed improvements on large datasets, as shown in experiments on TCGA data, and can be more easily interfaced with modern python-based data science tools.

**Availability and Implementation:** PyDESeq2 is released as an open-source software under the MIT license. The source code is available on GitHub at https://github.com/owkin/PyDESeq2.

## 1 Introduction

Bulk RNA sequencing (RNA-seq) is one of the most common molecular data modality used in biomedical research. Most RNA-seq datasets are used primarily for differential expression analysis (DEA) (Stark *et al*., 2019) which provides invaluable insight on the associations between the genes’ expression and a phenotype. Due to the inherent noise and statistical challenges present in RNA-seq data, DEA methods have become more sophisticated over the past decade, making them difficult to re-implement or port over new programming languages. In practice the community now relies primarily on a small handful of packages implementing state-of-the-art methods, among which `DESeq2` (Love *et al*., 2014).

While bioinformatics software is classically developed in R, a recent trend has seen the arrival of python software. Examples include the `scanpy` suite (Wolf *et al*., 2018) or the squidpy package (Palla *et al*., 2022) for single-cell and spatial RNA-seq, among others. This shift is motivated by several advantages of the python language: (1) the possibility to rely on well-maintained and efficient scientific computing packages such as `numPy` and `sciPy`, (2) a greater interoperability with machine learning and data science frameworks and (3) the potential to reach a wider audience, as python is one of the most popular programming languages (see, e.g., https://pypl.github.io/). Yet, to the best of our knowledge, there is currently no available python-native package for DEA with generalized linear models on bulk RNA-seq data.

A workaround consists in relying on python-to-R bindings, i.e. call R software and make back-and-forth data conversions from a python interface, using packages such as `rpy2` (https://rpy2.github.io/). However, this approach raises several issues: (1) it requires the user to install and maintain packages both in python and in R, which is cumbersome, (2) it creates computational overhead, as data is being converted and passed from one framework to the other and (3) it may lead to a loss of control for the user, as the options and subroutines of the original packages are only accessible through the binding layer.

In an effort to alleviate those issues and to benefit from the advantages offered by python-based software, we present `PyDESeq2`, a python implementation of the bulk RNA-seq DEA methodology introduced by Love *et al*. (2014) and implemented in the R package `DESeq2`.

## 2 Implementation

`PyDESeq2` implements the DEA methodology of Love *et al*. (2014), which briefly consists in modeling raw counts using a negative binomial distribution. Dispersion parameters are first estimated independently for each gene by fitting a negative binomial generalized linear model (GLM), and then shrunk towards a global trend curve. In turn, dispersions are used to fit gene-wise log-fold changes (LFC) between cohorts, and to perform Wald tests for differential expression.

### 2.1 Available features and code structure

For now, the features implemented in `PyDESeq2` correspond to default `DESeq2` settings. More precisely, it implements DEA for single-factor designs using Wald tests, and LFC shrinkage using the `apeGLM` prior (Zhu *et al*., 2019). Similarly to `DESeq2`, `PyDESeq2` is structured around two classes of objects: a `DeseqDataSet` class, handling data-modeling steps from normalization to LFC fitting, and a `DeseqStats` class for statistical tests and optional LFC shrinkage. To fit GLMs, we rely on the popular `scipy` (Virtanen *et al*., 2020) and `statsmodels` (Seabold and Perktold, 2010) python packages.

### 2.2 Comparison with DESeq2 on TCGA datasets

In Fig. 1 we compare the results of `PyDESeq2` and `DESeq2` on 8 datasets from The Cancer Genome Atlas (TCGA, https://www.cancer.gov/tcga). More precisely, we test differential expression between tissue samples corresponding to *advanced* vs. *non-advanced* tumor grades (as per TCGA’s clinical data), and focus on 4 criteria: retrieved genes, enriched pathways obtained with the `fgsea` package (Sergushichev, 2016), model likelihood, and speed.

**Figure 1:**
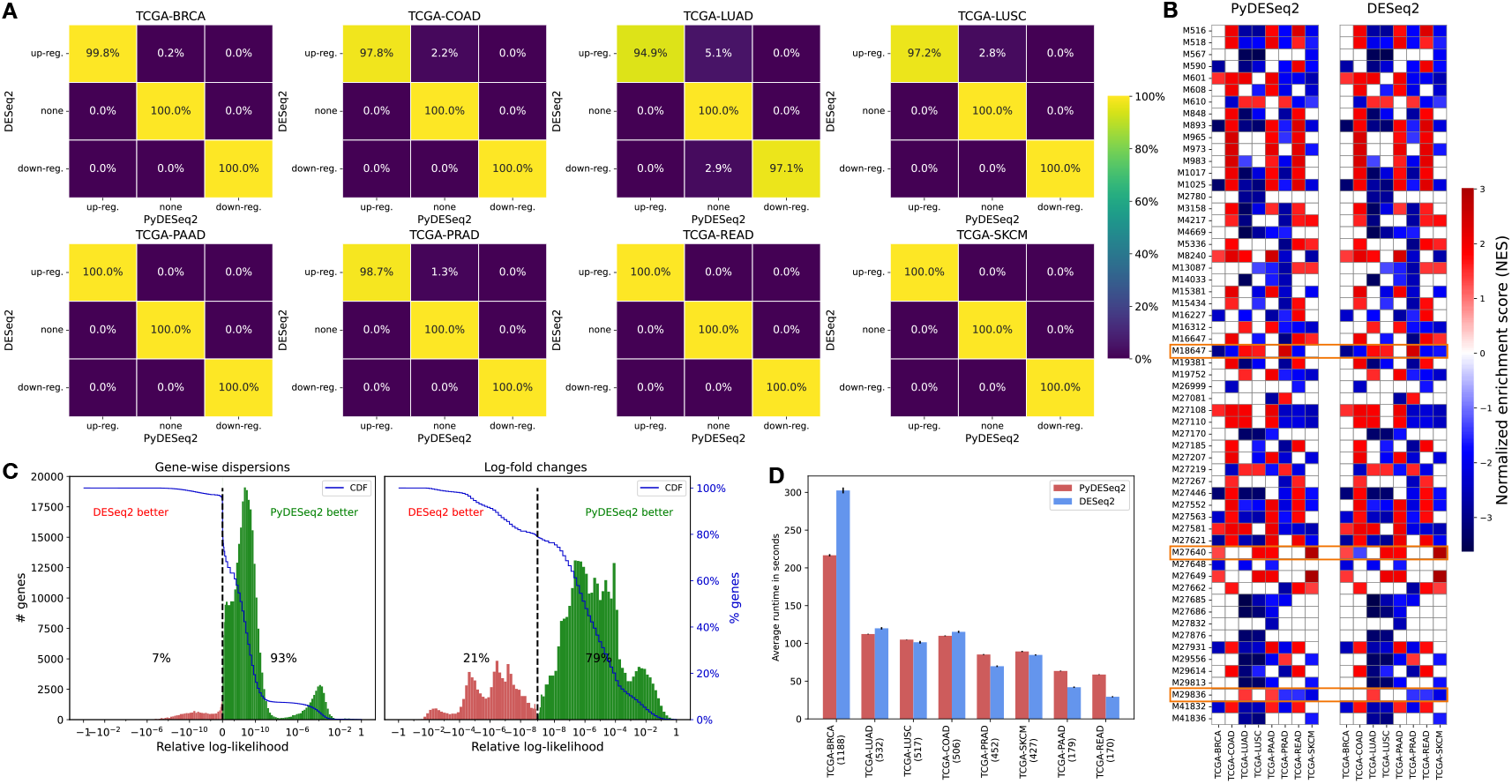
**(A)** Significantly differentially expressed genes (with padj ≤ 0.05 and |LFC| ≥ 2) according to `PyDESeq2` and `DESeq2`. **(B)** Significantly enriched pathways (padj ≤ 0.05) obtained with the fgsea package, using Wald statistics as gene-ranking metric. Only top 10 enriched pathways (according to adjusted p-value) of at least one cancer dataset are represented. If for a given cancer dataset, a pathway is not significantly enriched, the corresponding square is left blank. The 3 pathways which are considered significantly enriched in one implementation but not the other on a given TCGA dataset are highlighted by a surrounding box. **(C)**: Distribution of relative log-likelihoods 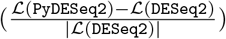, with corresponding cumulative distribution functions. **(D)** Time benchmark on an 8-core machine, averaged over 10 runs, using 8 threads for each package. Numbers between parenthesis correspond to dataset sample sizes. (A-D) We refer to the appendix for additional details on the experiments.

As can be seen from Fig. 1, `PyDESeq2` returns very similar sets of significant genes and pathways, while achieving better likelihood for dispersion and LFC parameters on a vast majority of genes, and at comparable speeds (higher for large cohorts, lower for small cohorts).

The data used in our experiments is publicly available on the TCGA Research Network website: https://portal.gdc.cancer.gov/. We refer to the supplementary material for additional details on the experiments.

### 2.3 Conclusion and future perspectives

In conclusion, `PyDESeq2` is a fast and reliable package for bulk RNA-seq DEA. By releasing this package, we hope to fill a gap in the python omics ecosystem, and to contribute to popularize the usage of modern data science python tools in gene expression analysis.

Finally, let us mention some of the features that we plan to implement in `PyDESeq2`. Future work includes adding support for multi-factor designs, and implementing the features induced by the `glmGamPoi` (Ahlmann-Eltze and Huber, 2020) option, such as using log-ratio tests and local median regression for the dispersions trend curve.

## Supporting information

Additional details

## Acknowledgements

The authors would like to thank Aura Moreno-Vega, Valérie Ducret and Quentin Bayard for fruitful discussions.

## Notes

https://github.com/owkin/PyDESeq2

### Competing Interest Statement

The authors are employees of Owkin, Inc.

## References

Ahlmann-Eltze, C. and Huber, W. (2020). glmGamPoi: fitting gamma-poisson generalized linear models on single cell count data. Bioinformatics, 36(24), 5701–5702.

Love, M. I. et al. (2014). Moderated estimation of fold change and dispersion for RNA-seq data with DESeq2. Genome biology, 15(12).

Palla, G. et al. (2022). Squidpy: a scalable framework for spatial omics analysis. Nature methods, 19(2), 171–178.

Seabold, S. and Perktold, J. (2010). statsmodels: Econometric and statistical modeling with python. In 9th Python in Science Conference.

Sergushichev, A. A. (2016). An algorithm for fast preranked gene set enrichment analysis using cumulative statistic calculation. bioRxiv.

Stark, R. et al. (2019). RNA sequencing: the teenage years. Nature Reviews Genetics, 20(11), 631–656.

Virtanen, P. et al. (2020). SciPy 1.0: Fundamental Algorithms for Scientific Computing in Python. Nature Methods, 17, 261–272.

Wolf, F. A. et al. (2018). SCANPY: large-scale single-cell gene expression data analysis. Genome biology, 19(1), 1–5.

Zhu, A. et al. (2019). Heavy-tailed prior distributions for sequence count data: removing the noise and preserving large differences. Bioinformatics, 35(12), 2084–2092.

