## Additional details for "PyDESeq2: a python package for bulk RNA-seq differential expression analysis"

### A Additional details on the experiments

#### A.1 PyDESeq2 and DESeq2 pipelines

For each dataset, we start with common row and column filtering steps. First, only samples whose tumor grade information (either *advanced* or *non-advanced*) information are kept. Then, only genes with a total sum of read counts over those samples are considered in the study.

We used the default settings of both `PyDESeq2` (with python 3.8) and `DESeq2` v 1.34.0 (corresponding to the "parametric" `fitType` for the latter), refitting Cooks outlier with a minimum number of replicates of 7, and performing an independent filtering step on the p-values to obtain ajdusted p-values with a threshold of 0.05 (note however that this threshold is not the default one in `DESeq2`). LFCs did not undergo a shrinking step.

In Fig. 1A, the genes are reported as significantly over (resp. under) expressed if the resulting adjusted p-value is under a 0.05 threshold, and if the corresponding log2-fold change is above (resp. under) a 2 threshold. Figures 3 to 10 display a detailed comparison of the retrieved genes for each dataset.

#### A.2 Gene set enrichment analysis

For GSEA, the results of the `PyDESeq2` and `DESeq2` pipelines were processed using a common R script based on the `fgsea` package (Sergushichev, 2016). This script uses Wald statistics as a gene-ranking metric, and sorts pathways based on the computed adjusted p-value. Pathways with an adjusted p-value under 0.05 were considered significantly enriched.

Enrichment was tested for pathways from the `C2:CP:REACTOME` collection of the Molecular Signatures Database (MSigDB, `http://www.gsea-msigdb.org/`).

An index of all pathways appearing in Fig. 1B is available in Table 1.

Detailed GSEA plots for the 3 pathways/dataset combinations with significant differences visible in Fig. 1B are represented in Fig. 2.

#### A.3 Time benchmark

The time benchmarks summarized in Fig. 1D were run on a dedicated GCP instance of the `n2-standard-8` type, with 8 virtual CPUs and 32GB of memory.

For each of the 8 datasets, the `PyDESeq2` and `DESeq2` pipelines were run using 8 cores, that is, setting `n_cpus` = 8 for `PyDESeq2` and `parallel` = `TRUE`, `BPPARAM` = `MulticoreParam`(8) in `DESeq2`.

Each pipeline was run 10 times. Fig. 1D represents the resulting average time using solid bars, and the standard deviation with error bars.

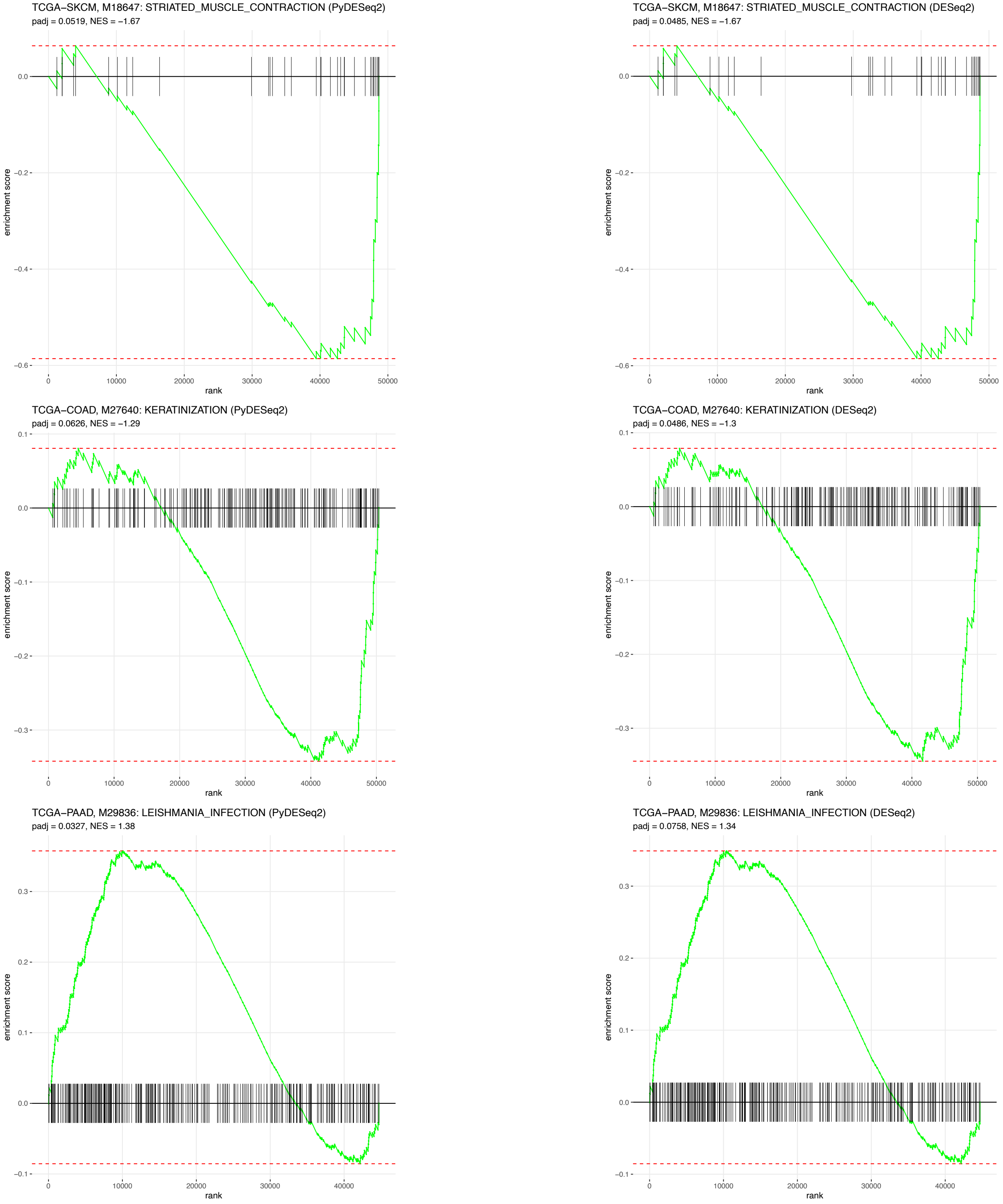


Figure 2: Zoom on the 3 pathway/dataset combinations from Fig. 1B for which `PyDESeq2` and `DESeq2` significantly differ. As one can see, in all 3 cases the GSEA plots and enrichment scores are nearly identical, but the adjusted p-value falls short of the padj $\leq 0.05$ for one implementation but not the other. This explains the differences observable in Fig. 1B (blank squares vs enriched pathways).

Table 1: Index of the pathways appearing in Fig. 1.

| ID | Pathway |
|---|---|
| M516 | THE CITRIC ACID TCA CYCLE AND RESPIRATORY ELECTRON TRANSPORT |
| M518 | ANTIGEN PROCESSING CROSS PRESENTATION |
| M567 | SRP DEPENDENT COTRANSLATIONAL PROTEIN TARGETING TO MEMBRANE |
| M590 | MITOCHONDRIAL PROTEIN IMPORT |
| M601 | ANTIGEN ACTIVATES B CELL RECEPTOR BCR LEADING TO GENERATION OF SECOND MESSENGERS |
| M608 | SIGNALING BY THE B CELL RECEPTOR BCR |
| M610 | EXTRACELLULAR MATRIX ORGANIZATION |
| M848 | MITOTIC G1 PHASE AND G1 S TRANSITION |
| M893 | RESPIRATORY ELECTRON TRANSPORT |
| M965 | INTERFERON GAMMA SIGNALING |
| M973 | INTERFERON ALPHA BETA SIGNALING |
| M983 | INTERFERON SIGNALING |
| M1017 | DNA REPLICATION |
| M1025 | RESPIRATORY ELECTRON TRANSPORT ATP SYNTHESIS BY CHEMIOSMOTIC COUPLING AND HEAT PRODUCTION BY UNCOUPLING PROTEINS |
| M2780 | SIGNALING BY ROBO RECEPTORS |
| M3158 | S PHASE |
| M4217 | MITOTIC PROMETAPHASE |
| M4669 | INFLUENZA INFECTION |
| M5336 | CELL CYCLE MITOTIC |
| M8240 | IMMUNOREGULATORY INTERACTIONS BETWEEN A LYMPHOID AND A NON LYMPHOID CELL |
| M13087 | PROCESSING OF CAPPED INTRON CONTAINING PRE MRNA |
| M14033 | MRNA SPLICING |
| M15381 | TCR SIGNALING |
| M15434 | DNA REPAIR |
| M16227 | CHOLESTEROL BIOSYNTHESIS |
| M16312 | CELL SURFACE INTERACTIONS AT THE VASCULAR WALL |
| M16647 | CELL CYCLE CHECKPOINTS |
| M18647 | STRIATED MUSCLE CONTRACTION |
| M19381 | G2 M CHECKPOINTS |
| M19752 | COMPLEMENT CASCADE |
| M26999 | COLLAGEN BIOSYNTHESIS AND MODIFYING ENZYMES |
| M27081 | METABOLISM OF STEROID HORMONES |
| M27108 | FCGR ACTIVATION |
| M27110 | ROLE OF PHOSPHOLIPIDS IN PHAGOCYTOSIS |
| M27170 | SELENOAMINO ACID METABOLISM |
| M27185 | MITOTIC METAPHASE AND ANAPHASE |
| M27207 | FCERI MEDIATED NF KB ACTIVATION |
| M27219 | ECM PROTEOGLYCANS |
| M27267 | TRANSCRIPTIONAL REGULATION BY TP53 |
| M27446 | MITOCHONDRIAL TRANSLATION |
| M27552 | TNFR2 NON CANONICAL NF KB PATHWAY |
| M27563 | DEFECTIVE CFTR CAUSES CYSTIC FIBROSIS |
| M27581 | CD22 MEDIATED BCR REGULATION |
| M27621 | COMPLEX I BIOGENESIS |
| M27640 | KERATINIZATION |
| M27648 | COPI MEDIATED ANTEROGRADE TRANSPORT |
| M27649 | FORMATION OF THE CORNIFIED ENVELOPE |
| M27662 | M PHASE |
| M27685 | RRNA PROCESSING |
| M27686 | EUKARYOTIC TRANSLATION INITIATION |
| M27832 | METABOLISM OF STEROIDS |
| M27876 | REGULATION OF EXPRESSION OF SLITS AND ROBOS |
| M27931 | NEGATIVE REGULATION OF NOTCH4 SIGNALING |
| M29556 | EUKARYOTIC TRANSLATION ELONGATION |
| M29614 | SEPARATION OF SISTER CHROMATIDS |
| M29813 | RESPONSE OF EIF2AK4 GCN2 TO AMINO ACID DEFICIENCY |
| M29836 | LEISHMANIA INFECTION |
| M41832 | REGULATION OF HMOX1 EXPRESSION AND ACTIVITY |
| M41836 | CELLULAR RESPONSE TO STARVATION |

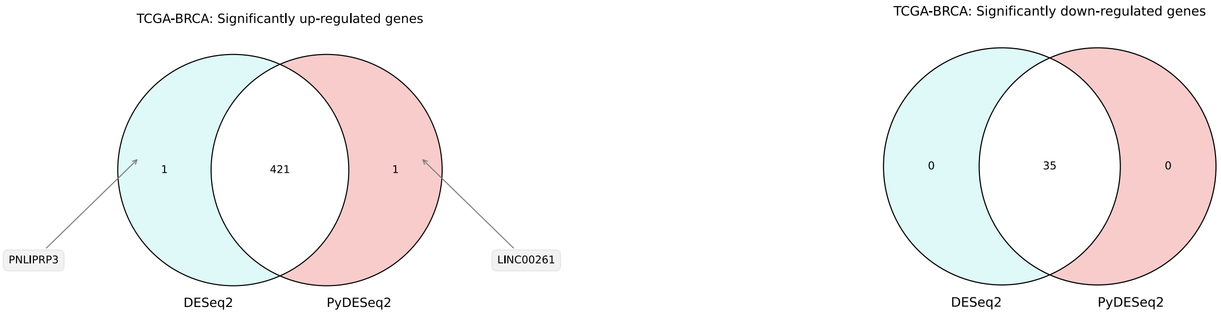


Figure 3: TCGA-BRCA: `PyDESeq2` and `DESeq2` retrieved significantly differentially expressed genes (padj $\leq$ 0.05 and $|\mathrm{LFC}| \geq 2$).

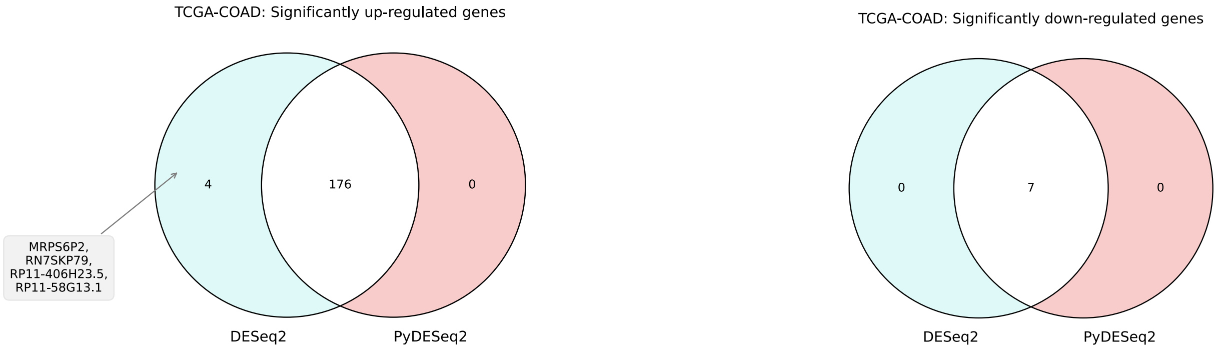


Figure 4: TCGA-COAD: `PyDESeq2` and `DESeq2` retrieved significantly differentially expressed genes (padj $\leq$ 0.05 and $|\mathrm{LFC}| \geq 2$).

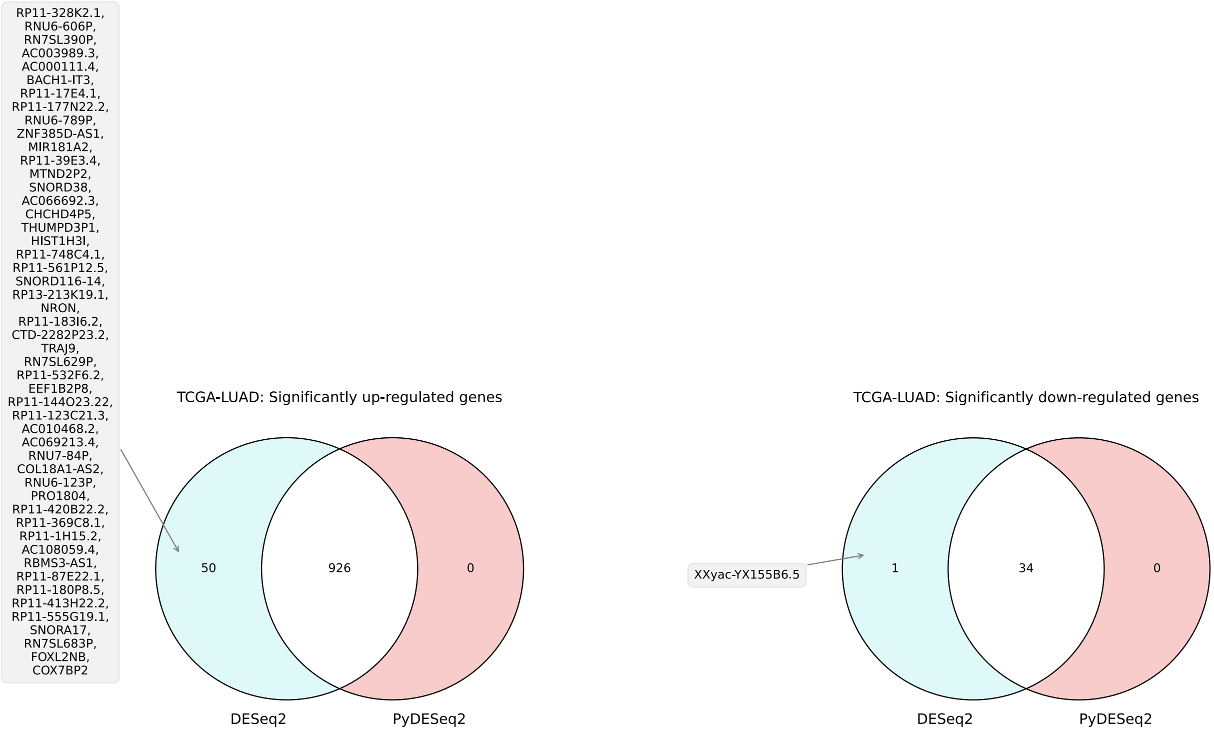


Figure 5: TCGA-LUAD: `PyDESeq2` and `DESeq2` retrieved significantly differentially expressed genes (padj $\leq$ 0.05 and $|\text{LFC}| \geq 2$).

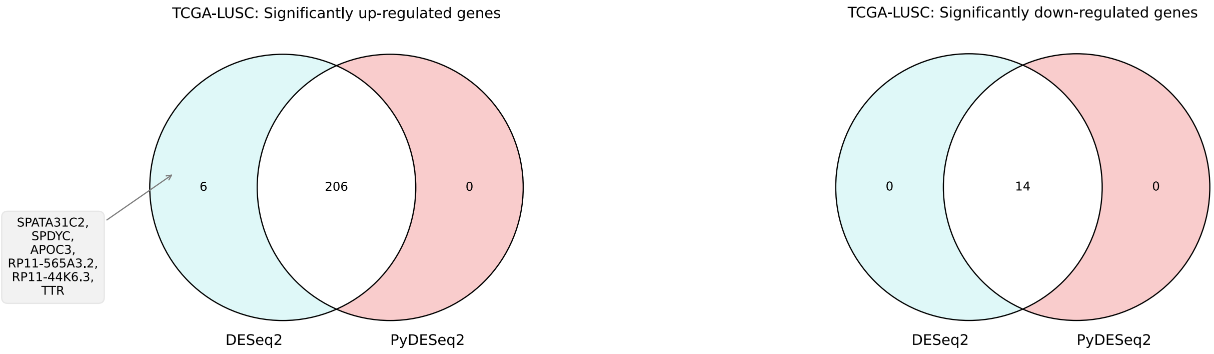


Figure 6: TCGA-LUSC: `PyDESeq2` and `DESeq2` retrieved significantly differentially expressed genes (padj $\leq$ 0.05 and $|\text{LFC}| \geq 2$).

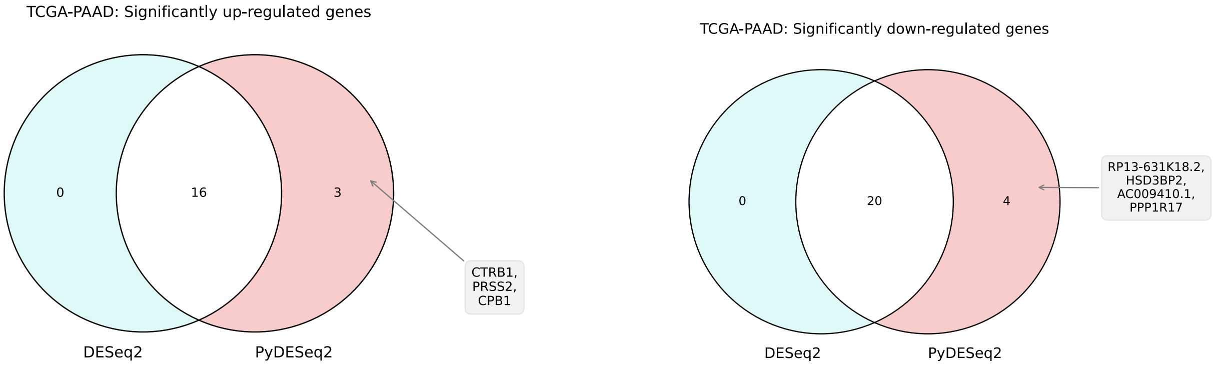


Figure 7: TCGA-PAAD: `PyDESeq2` and `DESeq2` retrieved significantly differentially expressed genes (padj $\leq$ 0.05 and $|\text{LFC}| \geq 2$).

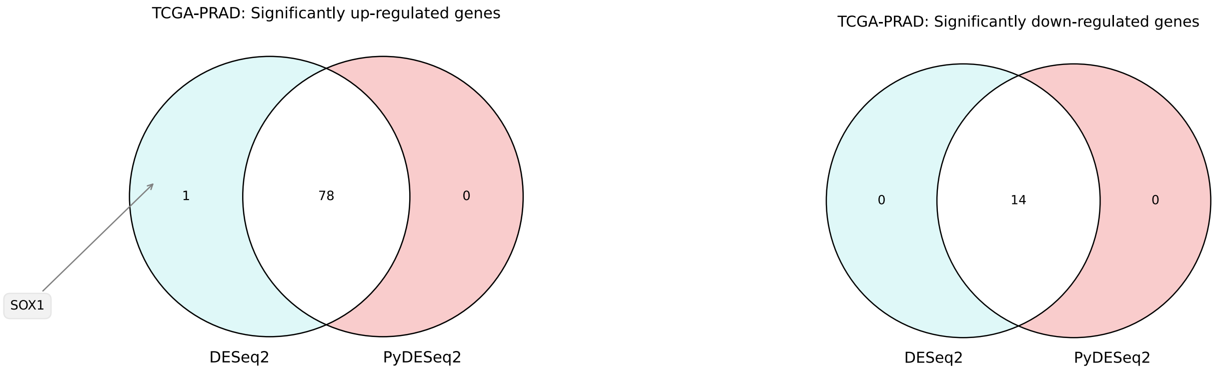


Figure 8: TCGA-PRAD: `PyDESeq2` and `DESeq2` retrieved significantly differentially expressed genes (padj $\leq$ 0.05 and $|\text{LFC}| \geq 2$).

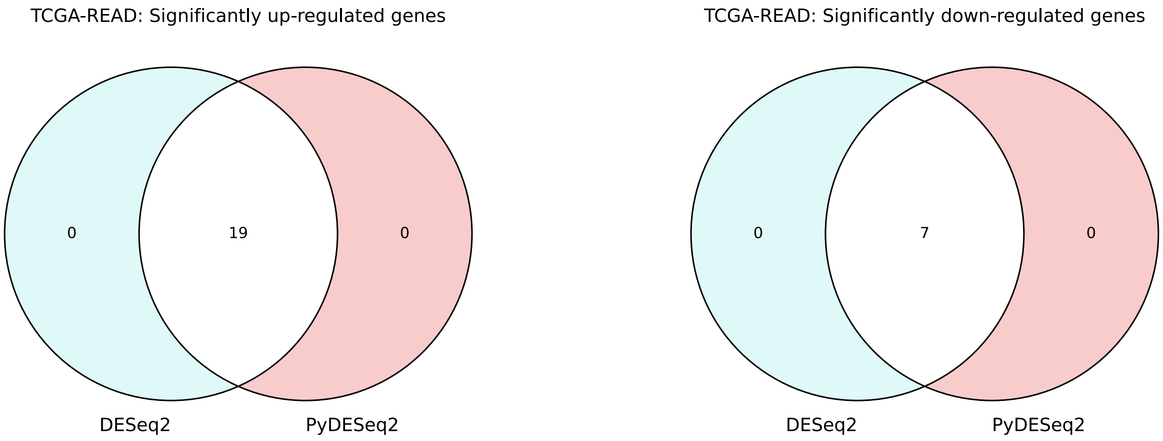


Figure 9: TCGA-READ: `PyDESeq2` and `DESeq2` retrieved significantly differentially expressed genes (padj $\leq$ 0.05 and $|\text{LFC}| \geq 2$).

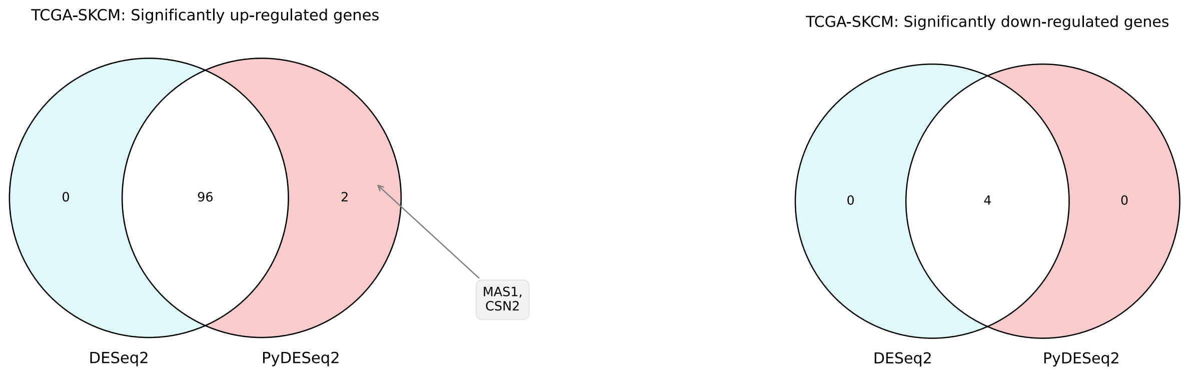


Figure 10: TCGA-SKCM: `PyDESeq2` and `DESeq2` retrieved significantly differentially expressed genes (padj $\leq 0.05$ and $|\text{LFC}| \geq 2$).
